# A major locus on chromosome 14 impacts developmental variation of Atlantic salmon smoltification

**DOI:** 10.1101/2025.10.16.682803

**Authors:** Marie Saitou, Domniki Manousi, Jayme van Dalum, Anja Striberny, Lars Grønvold, Cathrine Brekke, Solomon Antwi Boison, Jun Kwak, Arturo Vera Ponce de León, Bjarne Gjerde, Even Jørgensen, David Hazlerigg, Simen Rød Sandve

## Abstract

**Background:** Smoltification in anadromous Atlantic salmon is a complex developmental process involving physiological and cellular changes that enable freshwater fish to adapt to seawater. Central to this transformation is the functional transformation of the gill to manage osmoregulatory demands. While environmental cues like photoperiod are known to influence smolt development, the genetic architecture underlying smolt development — particularly related to gill physiology — remains poorly understood.

**Results:** A large-scale eQTL analysis across 3,000 Atlantic salmon subjected to three photoperiod regimes, identified over 45,000 significant SNP-gene expression associations in gill tissue. Notably, we discovered a 0.5 Mbp large trans-eQTL hotspot on chromosome 14. This ‘hub-locus’ was associated with expression of more than 2,000 genes across the genome which were significantly enriched for gill cell type specific markers. In addition we found that the hub-locus was associated with somatic growth. Our findings support a “local tissue effect” model, where cis-regulatory or protein sequence variants within the hub-locus modulate cell proliferation and differentiation of gill cell types.

**Conclusion:** This work advances our understanding of the genetic basis of smoltification in Atlantic salmon and provides a foundation for future studies using single-cell approaches to resolve cell-type specific mechanisms underlying genetic variation in smolt development.

## Introduction

Many species of fish undertake mass migrations linked to spatial and seasonal partitioning of feeding opportunities. In anadromous salmonids, migration of immature individuals from freshwaters to the sea allows exploitation of the marine environment to support rapid somatic growth, which is necessary for successful reproduction. The process of transforming from a freshwater resident to a seawater adapted migratory form is known as smoltification. Smoltification is a complex process involving many changes in physiological and behavioural traits, amongst which the reversal of osmoregulatory function from a water excreting, ion retaining mode to a water retaining ion excreting mode is of paramount importance (McCormick 2013).

The gill is the key organ through which smoltification related changes in osmo- and ion regulatory capacity take place (Evans *et al*. 2005). The high gas permeability of gill respiratory surfaces entails a concomitant high permeability to water and to solutes, and as a consequence gills contain specialised cells known as ionocytes (or mitochondria-rich cells), which control ion and water fluxes between the blood and the external water environment. This energy-expensive process, reliant on combined actions of ion transporting ATP-ases (principally sodium / potassium ATPase) and a range of ion cotransporters, leads to an osmoregulatory / respiratory compromise in gill function. Managing this osmo-respiratory compromise in freshwater and seawater environments requires radical adjustments in ionocyte function during smoltification, which manifest corresponding shifts in the distribution and cytological attributes of ionocytes (McCormick 2013).

In wild salmonids migration to sea generally takes place in a seasonally restricted spring time window. While the precise timing of migration in this window appears to depend on water levels and temperatures, which vary from year to year, the long term (i.e. annual) scheduling of migration is under photoperiodic control (i.e. a robust predictor of annual phase) (McCormick 2013). Indeed, sequential exposure to declining photoperiods in the autumn prior to smoltification followed by increasing photoperiods in the spring lead to photoperiodic history-dependent changes in pituitary hormone secretion (Strand *et al*. 2018) and gill gene expression profile (Iversen *et al*. 2020; Grønvold *et al*. 2024). These govern preparative changes in gill cytology and function months before seawater entry (Solbakken *et al*. 1994). Accordingly, in farmed salmonids, such as the Atlantic salmon (*Salmo salar*), artificial photoperiod manipulation has become a widely used component of the large-scale production of smoltified fish (smolts) (Ytrestøyl *et al*. 2023).

In addition to environmental cues, genetic variation also gives rise to variation in smolt development, including genotype and environment interaction. Khaw et al. (2021) identified significant heritability for a molecular gene expression phenotype associated with smolt development and development of seawater tolerance (Takvam *et al*. 2024), and recently Gjerde et al. (2025) demonstrated substantial heritability (h^2^=0.4 ± 0.06) for morphological smolt development characteristics. However, our understanding of the genetic architecture of smolt development is still meager, specifically related to key aspects of gill physiology, an essential aspect of the development of seawater tolerant smolts.

To address this knowledge gap, we designed a large-scale genetics experiment to identify genetic variation associated with smolt gill development under 3 different photoperiod treatment regimes. This led us to discover a ‘hub-locus’ on chromosome 14 which affects the expression of thousands of gill transcripts expressed from loci throughout the genome. These transcriptome-wide effects correlate with variation in growth phenotypes and represent a genetic locus at the core of developmental variation in smoltification of Atlantic salmon.

## Results

### Variation in smolt attributes under different lighting regimes

To study the impact of genetics and environmental (i.e. photoperiodic history) effects on smolt development we raised fish on 3 photoperiod regimes commonly used in the aquaculture industry (LD12:12, LD8:16, LL), which is known to produce different trajectories of smolt development **(Fig. 1A)**. As expected, we observed significant variation between- and within photoperiod treatment groups for smolt development phenotypes (previously described in (Grønvold *et al*. 2024; Gjerde *et al*. 2025)), e.g. body weight **(Fig. 1B)**, acute seawater phase mortality **(Fig. 1C)**.

**Figure 1.**
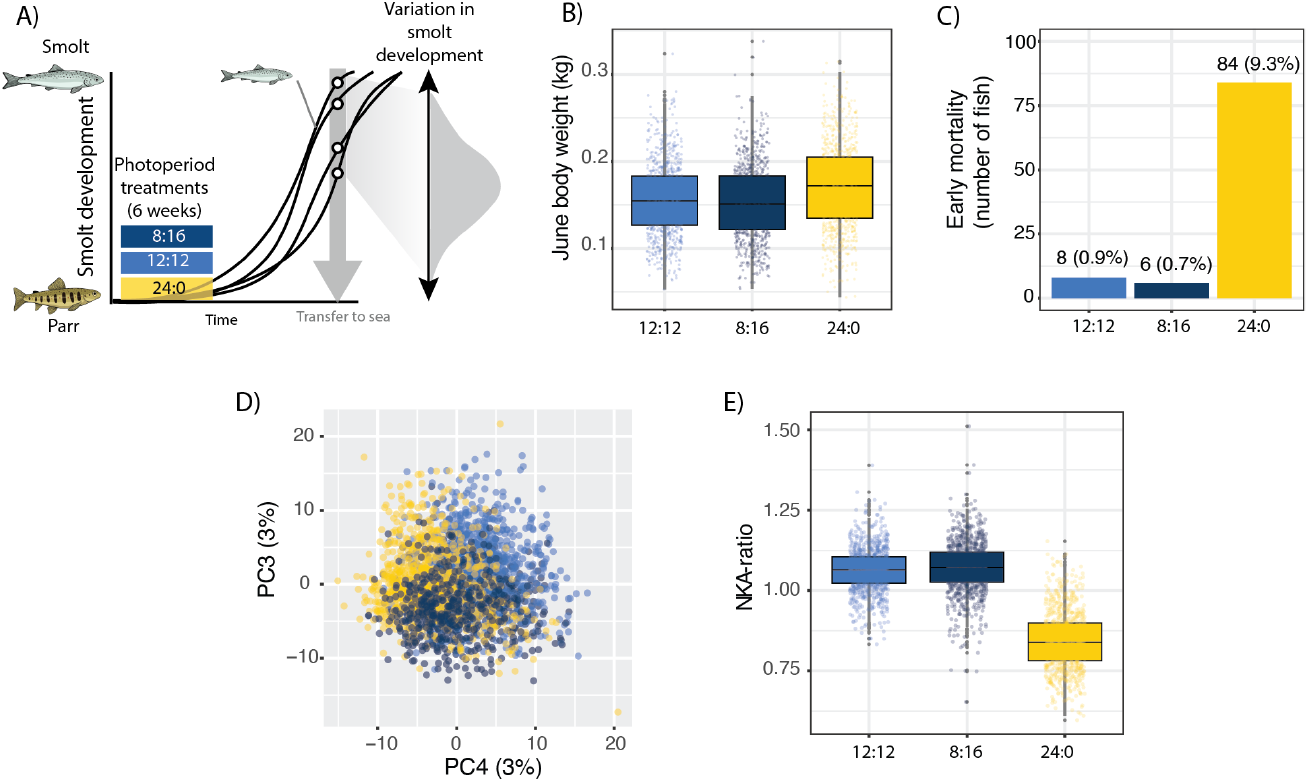
Experimental concept and variation in smolt development. A) The experimental design. Fish were taken through three smoltification protocols using different photoperiod regimes provoking variation in smolt physiology development within and between groups. Examples of theoretical smolt development trajectories for individual fish are indicated with lines. B) Body weight at time of sea transfer for the three photoperiod treatment groups. C) Early mortality after sea water transfer for the three photoperiod treatment groups. D) PCA scatterplot of transcriptome variation across 2806 fish. The principal components 3 and 4 showed signatures of photoperiod group effects and are visualized. E) Normalized gene expression levels (transcripts per million) for the smolt gill development gene expression marker NKA-ratio: the ratio of *nka*-*α1b vs nka*-*α1a* (LOC106575572/LOC106602157).

Development of sea water adapted gill physiology is a critical aspect of smolt development, however direct measurement of the variation in ‘gill function’ across thousands of samples is not feasible. We therefore generated RNA-seq from small gill biopsies from all fish and used the gill transcriptome as a proxy for smolt gill development. To assess overall RNA-seq data quality and tank reproducibility for the molecular traits we first undertook PCA-based visual explorative analyses of the RNA-seq data.

The PCA-based visual explorative analyses of the RNA-seq data from 2806 fish revealed a weak but visually detectable genome-wide transcriptome effect of photoperiodic treatment and identified large within- -group variation in gill transcriptome phenotypes **(Fig. 1D)**. However, zooming in on specific smolt-gill developmental genes revealed large within- and between-group gene expression variation, in line with previous studies on smolt gill development and our expectations from the experimental design **(Fig. 1E)**.

### Identification of a genetic ‘hub locus’ shaping gill development

We used an eQTL approach to identify 45,543 significant associations between SNP genotype variation (total SNPs = 415,000; 366,437 imputed and 48,563 SNPs from the genotyping array) and the expression level of 16,314 genes in the gill (**Fig. 2A**). Approximately 65% (29,419 / 45,543) of these associations were detectable in all photoperiod regimes, and more eQTLs were identified in 8:16 and 24:0 fish compared to 12:12 fish (**Fig. 2B**).

**Figure 2.**
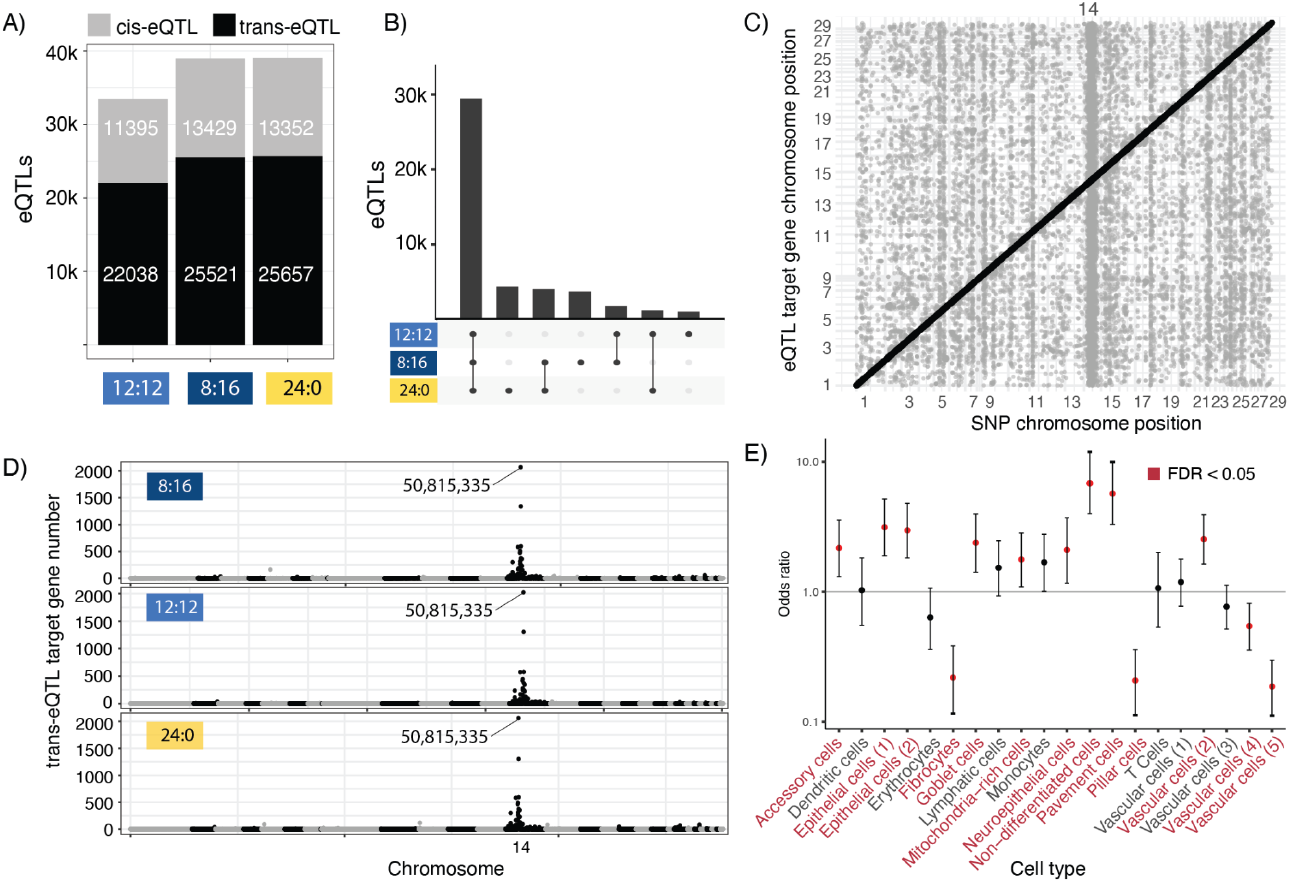
Gene expression phenotypes and eQTL results overview. A) Total number of eQTL associations. B) Shared and unique eQTL associations across light regimes. C) Dotplot of eQTL SNP position against target gene position. Black = cis-eQTLs, grey = trans-eQTLs. D) Chromosomal distribution of trans-eQTL targets gene numbers across chromosomes for each light regime. E) Enrichment analysis for gill cell type specific genes across the three hub-SNP genotypes. Positive enrichment can be interpreted as cell types that are more prevalent in the fish with minor allele genotype.

Subdividing eQTL associations into cis- (target gene close to SNP) and trans- (target gene >500kb from SNP) associations, we found about twice as many trans-compared to cis-eQTLs in each of the photoperiod regimes **(Fig. 2A)**. Visualizing the genomic coordinates of eQTL associations revealed several trans-association ‘hotspots’ in the genome (seen as vertical stripe patterns in **Fig. 2C**), and chromosome 14 stood out as an extreme outlier with respect to trans-eQTL signals (**Fig. 2C**), harbouring over 30% (11,857) of all trans-eQTL associations in the genome. Further investigation into the genetic source of this chromosome 14 effect identified a single genomic region (hereafter referred to as the ‘hub-locus’) centred at approximately 50.8 Mbp (SNP chr14_50815335, **Fig. 2D**). The hub-SNP had a minor variant with moderate frequency (0.25) and was associated with variation in expression levels of 2,075 unique target genes across all chromosomes **(Fig. 2C)**. Comparing hub-SNP trans-effects across photoperiod regimes showed that the hub-SNP targets were highly consistent with 96% of all significant hub-locus eQTL associations shared across all three photoperiod regimes.

To better understand the molecular underpinnings of the hub-locus we tested if the hub-locus target genes were associated with particular cell types or enriched for particular functions. Using a previously published sn-RNA dataset from Atlantic salmon gills (West *et al*. 2021), we found that the hub-locus target genes were enriched in cell type-specific markers from 16 gill cell types (Fisher test FDR<0.05, **Fig. 2E**). Significant enrichment signals (FDR<0.05) were observed for cell types such as mitochondria-rich cells, accessory cells, pavement cells, vascular cells, and non-differentiated cells, which are all known to change abundance through smolt development (West *et al*. 2021). A GO-enrichment test for the eQTL targets did not reveal any functional enrichment signals of notice.

### The hub-locus influences somatic growth

To better understand how the hub-locus impacts variation in smolt development we performed in-depth analyses of the hub-locus effect. We first characterized the hub-SNP eQTL photoperiod interaction effects and effect sizes. Compared to all trans-eQTLs (**Fig. 3A**), the hub-locus had half the amount of photoperiod interactions (2.5% vs 11.5%, **Fig. 3A, B**). Furthermore, the hub-locus trans-eQTLs generally had larger regression slope values (in absolute terms) compared to other trans-eQTLs **(Fig. 3C)**, but the differences in slopes (i.e. the magnitude of photoperiod interactions) were similar for hub-eQTLs and other eQTLs (**Fig. 3D**).

**Figure 3.**
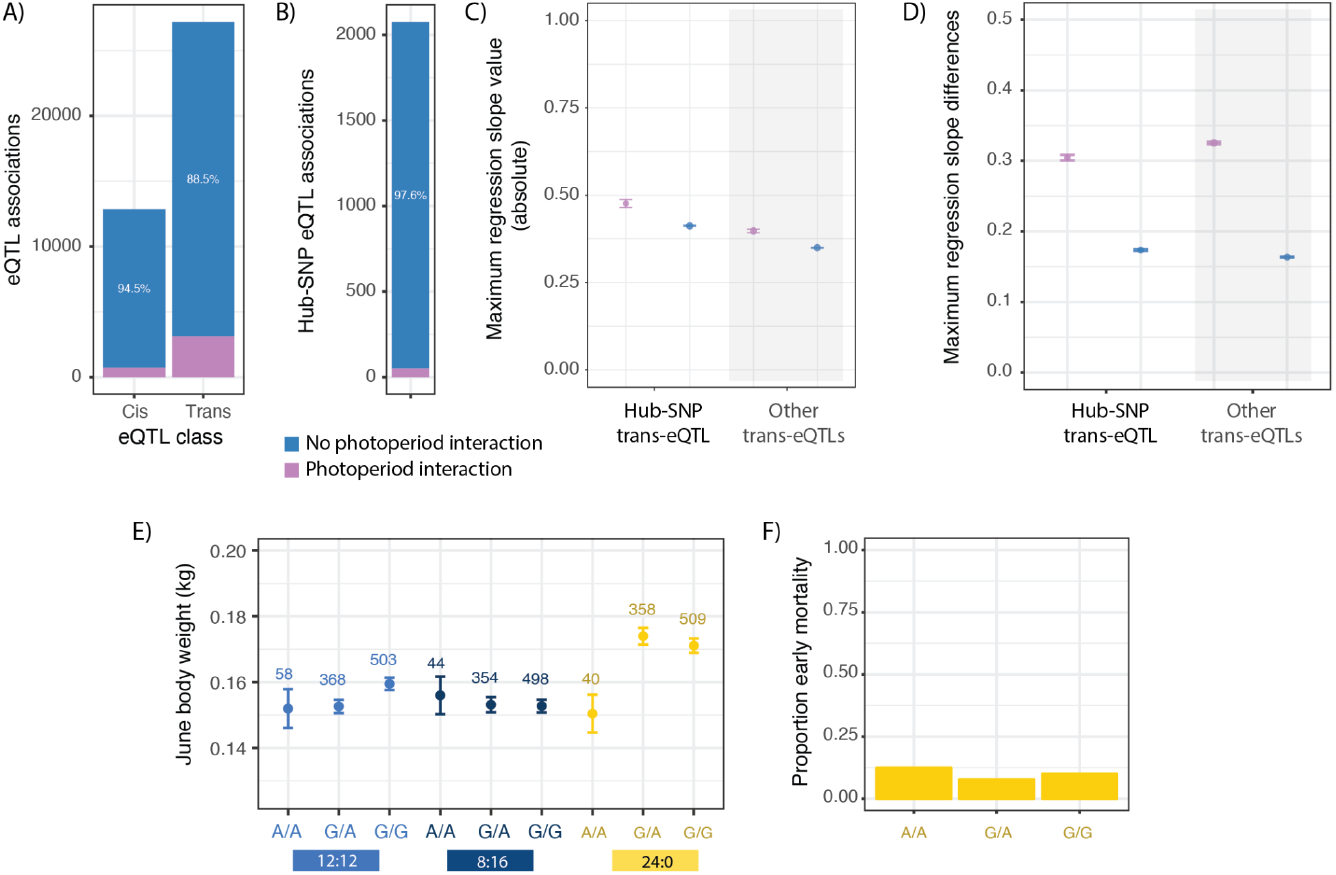
Interactions between photoperiod history and genetics on eQTLs. A) Number and proportion of cis- and trans-eQTLs with significant photoperiod history interaction effects. B) Number and proportion of hub-locus eQTLs with significant photoperiod history interaction effects. C) Maximum absolute value of regression slopes for trans-eQTL associations across the three photoperiodic treatments. D) Maximum difference in regression slope values across photoperiod treatments for trans-eQTL associations. E) June body weight for the three hub-SNP genotypes across the three photoperiodic treatments. F) Proportion of fish with early mortality for hub-SNP genotypes in the 24:0 treatment group.

Next we examined the associations between the hub-locus genotype and variation in body weight and sea water mortality, two aspects of smolt development known to vary extensively between photoperiod treatments **(Gjerde et al. 2025)** (**Fig. 1B, C)**. The 24:0 treatment group displayed significant variation in growth across the three genotypes for the hub-SNP (ANOVA *p*=0.015; **Fig 3E**), with fish homozygous for the hub-SNP minor allele having approximately 14% lower body mass. Early sea water mortality on the other hand, showed no clear association with hub-SNP genotype (Fisher exact test *p*-value = 0.38, **Fig. 3F**). This indicates that the hub-locus has a large photoperiod-independent effect on bulk tissue gill transcriptome variation and at the same time impacts somatic growth.

### Dissection of the hub-locus genetic effects

To dissect the genetic mechanism underlying the hub-locus effect on smolt developmental variation, we analysed linkage disequilibrium (LD)-structures and genomic variation present around the hub-region in the context of gene annotations **(Fig. 4A)**. LD analysis revealed an approximately 0.5 Mbp region with elevated LD surrounding the hub-SNP (R^2^ >0.5) and therefore low ability to confidently distinguish causal candidate variants. Nevertheless, the local LD structure surrounding the hub-locus plateaued over two genes (**Fig 4B**); PAK interacting protein (PAK1IP1, ENSSSAG00000004575) and uridylate-specific endoribonuclease C (ENDO-U, ENSSSAG00000004590). Prediction of variant effects for SNPs overlapping these genes did not identify good candidate causal variants classified as ‘high’ impact (i.e. missense) (Supplementary table S1), however this does not exclude that one or more of these variants could potentially impact the regulation of these loci.

**Figure 4.**
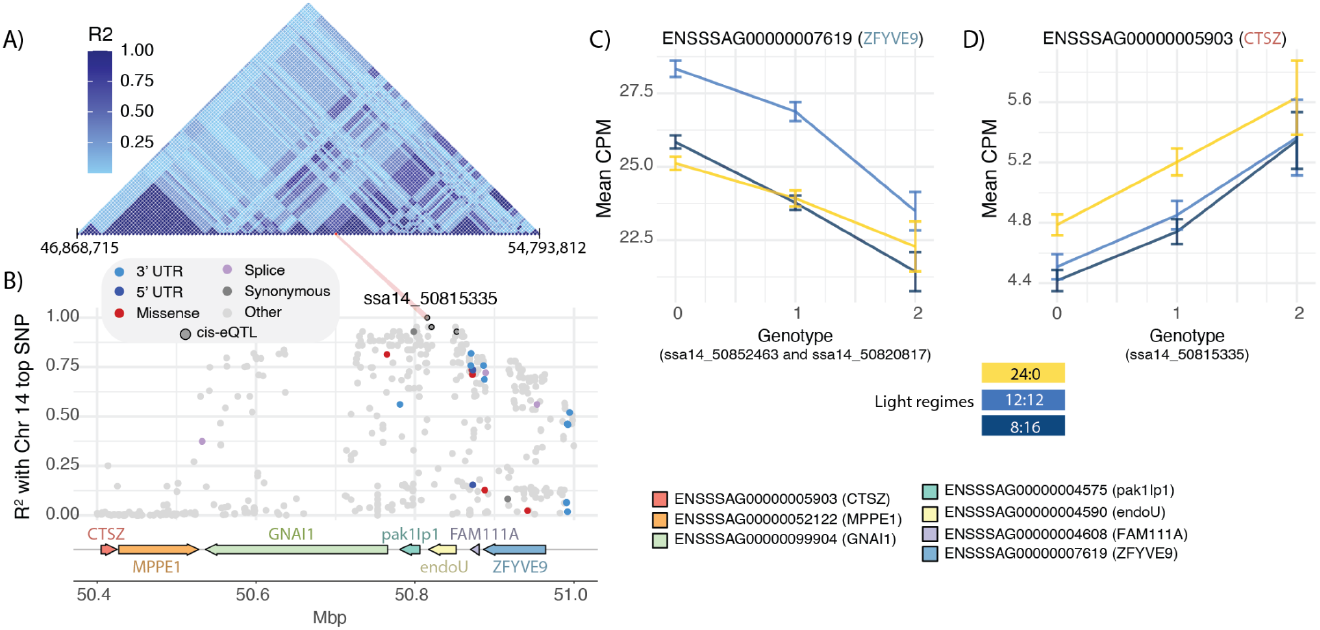
Dissection of the hub-locus genetic variation. A) LD-plot of SNP-markers flanking the hub-locus genomic region. B) Zoom-in on hub-SNP region showing SNP-markers (including imputed variants), their R2 to the hub-SNP, and protein coding gene models. Dot colour highlights variant prediction effect class and if a SNP is a cis-eQTL for a gene in the region and has R2<0.9 to the hub-SNP. C, D) Gene expression phenotypes for cis-eQTL associations between SNP-variants and genes in the hub-region.

To address this question we therefore identified cis-eQTLs (FDR<0.05) in very high LD (0.9) with the hub-SNP which also were associated with variation in gene expression levels for genes within the hub-locus region (Fig. 4B). Three SNPs matched these criteria, targeting the genes ENSSSAG00000007619 and ENSSSAG00000005903 (**Fig. 4C, D**). The two genes were relatively lowly expressed and the variants were associated with ∼12-15% change in transcription levels between the two homozygous genotypes. ENSSSAG00000007619 is orthologous to a gene annotated as ZFYVE9 in brown trout according to salmobase.org, and the human ortholog of this gene is associated with regulation of TGF-beta signalling according to genecards.org. The other gene (ENSSSAG00000005903) is according to the NCBI annotation of the Atlantic salmon genome encoding a protein similar to cathepsin Z (CTSZ).

Limiting detection of causal genetic variants to SNPs can compromise attempts to identify causal variation for complex traits. Hence, we used Oxford Nanopore long-read sequencing technologies to obtain genotype information from two homozygous individuals for the hub-locus and use simple as well as complex (structural) genetic variants to search for potentially causal elements. To further refine our findings, we predicted the functional effect of complex variants on protein function. In total, 10,160 SVs detected on chromosome 14, of which 567 were homozygous in at least one of the two individuals and had a high predicted impact on protein function (Supplementary table 1). Two high-impact variants were deletions predicted to cause feature truncation in PAK1IP1 and ENDO-U. The homozygous genotype for the hub-locus minor variant carried a homozygous 99 bp deletion (Sniffles2.DEL.11D6M7) at 50,787,668–50,787,766, affecting coding exon 10 of PAK1IP1. The homozygous genotype for the hub-locus major variant carried a homozygous 91 bp deletion (Sniffles2.DEL.11EBM7) of 50,823,338–50,823,428, affecting coding exon 9 in ENDO-U.

## Discussion

In this study we identify a genomic region (the hub-locus: ssa14_50815335) on chromosome 14, associated with gill transcriptome variation impacting >2000 genes (**Fig. 2**). This type of trans-eQTL ‘hotspot’, impacting thousands of genes, has previously been reported in yeast (Albert *et al*. 2018; Renganaath and Albert 2025) where it is known to be linked to complex trait variation such as growth (Renganaath and Albert 2025). Similarly, we find that the hub-locus trans-eQTL was also associated with somatic growth in Atlantic salmon (**Fig. 3**). Although the exact genetic mechanisms underlying the hub-locus effect on smolt developmental variation remains unclear, our study provides some insights into possible mechanistic models.

Firstly, eQTL signals from bulk tissue RNA-seq data could either arise from genetic variation impacting RNA-abundance within a nucleus or alternatively through a correlation between a genotype and the cellular composition of a tissue sample (Zhang and Zhao 2023). Our study is based on a single time point in a developmental gradient and reveals significant enrichment of many cell type-specific markers among the target genes of the trans-eQTL hub-locus (**Fig. 2E**). We therefore argue that the hub-locus trans-eQTL signal is, at least partly, an effect of variation in cell type composition between genotypes, as a result of smolt gill-cell developmental programs out of sync, or at different trajectories.

But where does the hub-locus effect originate, and how does it translate to variation in the gill cell population structure? One possibility is that the hub-locus exerts its direct effect at a peripheral tissue regulating endocrine signals as variation in endocrine profiles could easily give rise to developmental variation in gill cell composition and body size (McCormick 2013). One prediction from this model is that the hub-locus should contain genes with key roles in endocrine regulation and signalling, which possibly also is reflected in brain biased gene expression profiles (note that under this model these genes do not need to have strong cis-eQTL signals in gill tissue). Two of these genes within the hub-locus, GNAL (annotated as guanine nucleotide-binding protein G(olf) subunit alpha in NCBI) and MPPE1 (annotated as metallophosphoesterase 1-like in other salmonid orthologs) (Fig. 4B), had clear brain-biased expression according to tissue expression profiles in salmobase.org. However, these genes have no known roles in endocrine regulation as far as we are aware of, which does not align with the ‘peripheral endocrine model’.

Another scenario is that the hub-locus exerts its direct effect in many cell types present in the gill through creating variation in these cells’ ability to receive, sense or respond to external signals that modify cell-proliferation or -differentiation (‘local tissue effect model’). Under this model the causal variant could be a regulatory mutation (likely a cis-eQTL variant within the hub-locus) or a coding sequence variant that impacts the function of one of the genes in the hub-locus. We found three cis-eQTL variants in very high LD with the hub-SNP. The hub-SNP itself was a cis-eQTL impacting CTSZ (**Fig 4D**). This gene is known to modulate cell proliferation and migration (Tao *et al*. 2025), with implications for immune cells in fish (Cai *et al*. 2019), and is found to be differentially expressed in gill tissues from smolts at different developmental stages (Morro *et al*. 2019). Two other putative causal cis-eQTL are found at ssa14_50852463 and ssa14_50820817 associated with the expression levels of gene ZFYVE9. In vertebrates, including fish, this gene participates in the transforming growth factor beta (TGF-beta) pathway (Tsukazaki *et al*. 1998) and plays a role in the formation of new tissue (Bensimon-Brito *et al*. 2020) and immune cell regulation (Maehr *et al*. 2012; Zhang *et al*. 2023). All these cis-eQTLs have, however, rather small effects on their target genes (**Fig. 4C, D**), questioning their potential to exert large effects on transcriptomes and complex traits. Yet, apparently small eQTL effect sizes from bulk tissue RNA-seq is sometimes due to an ‘averaging out’ effect across many cell types (Zhang and Zhao 2023), which only can be resolved using single cell-based approaches.

Using long-read sequencing data from fish with homozygous hub-locus genotypes we also found structural genetic variation with a potential impact on protein function of the two genes flanking the hub-SNP, PAK1IP1 and Endo-U (Fig. 4B). These two genes also have the potential to drive large scale gene expression variation. The downstream flanking gene (ENSSSAG00000004590, Endo-U) encodes RNAse enzymes which are found in viruses, bacteria and eukaryotes, but are generally poorly characterized(Malard *et al*. 2025). In mammals and frogs, orthologs of Endo-U have been shown to direct large-scale degradation of RNAs(Laneve *et al*. 2003; Schwarz and Blower 2014; Malard *et al*. 2025). The upstream flanking gene to the hub-SNP (ENSSSAG00000004575, PAK1IP1) is in humans involved in cell cycle regulation and cell proliferation (Yu *et al*. 2011; Panoutsopoulos *et al*. 2020), with potentially large effects on gill cell population development.

In conclusion, we have identified a large effect variant on chromosome 14 associated with developmental variation of the gill physiology and -growth during the smoltification of the fish. LD analysis pinpoints the association signal to a fairly small region (∼0.5 Mbp) containing multiple genes. We hypothesise a local tissue effect model whereby a casual variant (cis-regulatory or structural) within the hub-locus has a direct effect on gill cell proliferation and differentiation. This hypothesis can be tested by contrasting single cell RNA-seq data from various tissues across hub-locus genotypes.

## Methods

### Ethical statement

The experiment adhered to EU regulations for the protection of animals used in research (Directive 2010/63/EU). All necessary precautions were taken to reduce pain and discomfort. The Norwegian Food and Safety Authority approved the experiment (FOTS ID: 25658).

### Fish samples

Here follows a brief description of the fish rearing. A detailed description is published in Gjerde et al. (2025). We reared approximately 3000 Atlantic salmon (*Salmo salar*) from fertilized eggs to smolts over a period of 1 year and 3 months. The fish were divided into three groups subjected to different photoperiodic regimes. A ‘24:0’ group was maintained under continuous light (e.g. 24 hour light per day) from hatching until end of smolt development. The two other groups were subjected to a 6 week period of shorter days when they reached approximately 50g. The ‘12:12’ group were kept on 12 hours light / 12 hours dark and the ‘8:16’ group were kept on 8 hours light / 16 hours dark. Following this short photoperiod exposure 12:12 and 8:16 fish were transitioned back to continuous light for 8 weeks. Gill tissue biopsies for RNA-seq were collected from all fish in all groups within the same week, at the end of the smolt production protocol.

### SNP-chip genotyping

All gill samples were genotyped using a 66K SNP Affymetrix array [Salmow01]. Physical array coordinates were re-assigned to the newer Atlantic salmon genome reference (GenBank accession: GCA_905237065.2) and variants were quality-filtered using PLINK v1.9 (Chang *et al*. 2015) using the following thresholds: HWE p-value < 1E-6, MAF < 0.01. Samples with more than 10% of their genotypes missing were removed, and variants were removed if there were missing genotype calls in more than 3% of the fish. A total of 2970 fish and 48,563 SNP were retained after quality filtering.

### Whole genome sequencing

We sequenced 112 parent fish of the smolt population using short read illumina technology. Read mapping to the reference genome (GCA_905237065.2) was done using BWA (Li and Durbin 2009). Variant calling was done using the GATK pipeline (Van der Auwera *et al*. 2013). Quality filtering of SNP variants was done with PLINK v1.9 (Chang *et al*. 2015) based on Hardy-Weinberg Equilibrium (HWE p-value < 1E-8), minor allele frequency (MAF < 0.01), and genotype missingness thresholds per individual and SNP variant (10% and 5%, respectively).

### Genotype imputation

To improve the SNP coverage we performed genotype imputation on the smolt population using the parent genotypes from the whole genome sequencing data. Imputation was done using Beagle 5.2 software (Browning *et al*. 2021). In order to identify SNP with high inference accuracy, a five-fold cross-validation method was applied by dividing the parent fish into five groups. Imputed genotypes were masked in 20% of the individuals and then imputed using the remaining 80%. Imputation accuracy was assessed as the square of Pearson’s correlation coefficient (R^2^) between the true and imputed genotypes, averaged across the five validation results. This process resulted in 412,069 SNPs with an R^2^ > 0.80 which were further filtered using HWE (HWE p-value < 1E-8) using PLINK v1.9.

### RNA-sequencing

For a detailed description of RNA-seq data generation and expression quantification see Grønvold et al. (Grønvold et al. 2024). Briefly, gill biopsies were flash frozen, and RNA was extracted using the RNeasy Fibrous Tissue Mini kit (QIAGEN). Libraries were prepared and sequenced with 2×50 bp paired-end Illumina sequencing. Adaptor sequences were removed using fastp version 0.23.2 (Chen *et al*. 2018) and transcript quantification were conducted with Salmon version 1.1.0 (Patro *et al*. 2017) and the Atlantic salmon transcriptome annotation from the Ssal_v3.1 genome assembly (Ssal_v3.1, GCA_905237065.2) from Ensembl. We applied the selective mapping mode using a transcriptome index file as suggested by (Srivastava *et al*. 2020) . The --keepDuplicates and --gcBias options were used to address high sequence similarity between duplicated genes resulting from the salmonid genome duplication (Lien et al. 2016) and to correct for fragment-level GC biases, respectively. Gene-level mRNA expression was calculated by summing raw read counts and normalizing them into transcripts per million (TPM) using the R package tximport (Soneson *et al*. 2015).

### eQTL analysis

We first used QTLTools (Delaneau *et al*. 2017) to detect eQTLs for each light condition, regressing out covariates including father, mother, sex, tank, weight, body length, condition factor, skin colouration, and ten genomic principal components to control the genetic relatedness and batch information. Significant eQTL associations (FDR< 0.05) were defined as cis-eQTLs if the SNP-gene distance <5Mb and as trans-eQTL if SNP-gene distance > 5Mb or SNPs and genes were located on different chromosomes.

To assess the interaction between genotype and light treatment conditions on gene expression levels, we performed the following analyses. From the complete set of gene expression profiles and genotype data, we extracted gene-SNP pairs that showed a significant eQTL effect (FDR ≤ 0.05) in at least one of the three groups (12:12, 8:16, or LL).

We used the following linear model:

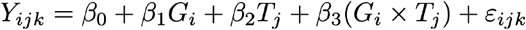

where *Y*_*ijk*_ is the expression level of gene *i* under photoperiod regime *j* for individual *k. β*_1_*G*_*i*_ is the fixed effect of genotype; . *G*_*i*_ denotes the genotype dosage (0, 1, or 2), *T*_*i*_ is the fixed effect of l photoperiod regime, and *β*_3_(*G*_*i*_ *× T*_*i*_) : is the Interaction effect between genotype and treatment; (*G*_*i*_) depends on the photoperiod regime (*T*_*i*_).

*ε*_*ijk*_ is residual error term for gene *i* in individual *k* under treatment *j* . The model was implemented using ordinary least squares regression.

### Long read sequencing and structural variation calling of two hub-SNP homozygous fish

Two Atlantic salmon individuals with homozygous genotypes for reference and alternative allele of the hub-locus SNP variant (chr14_50815335) were selected from the breeding nucleus of an aquaculture company (MOWI). Extraction of DNA for these samples followed standard protocols and genomic material was sequenced using Oxford Nanopore (ONT) sequencing technologies on the PromethION sequencing platform.

Raw Sequencing reads were quality filtered using Filtlong v0.2.1 to remove the worst 10% of reads as well as reads with length shorter than 4000 bp. Filtered output was then aligned against the Atlantic salmon genome (Ssal_v3.1) using minimap2 v2.26-r1175 (Li 2021), and structural variation (SV) calling was conducted individually for each fish using Sniffles2 v2.2 (Sedlazeck et al. 2018). Discovered SVs were then merged together using the latter software to create a long-read based SV catalog. From the produced catalog, elements overlapping chromosome 14 were filtered for further analysis using PLINK2 (Chang et al. 2015). Finally, the functional impact of filtered variants was predicted using the Ensembl Variant Effect Predictor and the Atlantic salmon functional annotation (Ssal_v3.1).

## Data and code availability

The illumina sequencing data from RNAseq and whole genome resequencing is available in the European Nucleotide Archive under the project PRJEB47441. The code and output from bioinformatics pipelines used to generate all the figures and downstream analyses of data in R is available at https://gitlab.com/sandve-lab/smolt_photogenetics.

## Acknowledgements

This study was supported by the Norwegian Seafood Research Fund (FHF) through project 901589 and Norwegian Research Council (NRC) through project 343129.

## Author contributions

SRS and DH conceived the study and provided funding. AS, JVD, EJ, DH planned and executed the fish rearing experiments and sampling. AS, SRS, SB, and DH organized the data generation. MS, JVD, LG, DM, SRS, CB, JK, APDL, BG and SB contributed to analyses of the data. SRS, DH, and MS drafted the manuscript. All co-authors took part in manuscript revisions.

## Notes

### Competing Interest Statement

The authors have declared no competing interest.

## Bibliography

Albert F. W., J. S. Bloom, J. Siegel, L. Day, and L. Kruglyak, 2018 Genetics of trans-regulatory variation in gene expression. eLife 7. 10.7554/eLife.35471

Bensimon-Brito A., S. Ramkumar, G. L. M. Boezio, S. Guenther, C. Kuenne, et al., 2020 TGF-β Signaling Promotes Tissue Formation during Cardiac Valve Regeneration in Adult Zebrafish. Dev. Cell 52: 9-20.e7. 10.1016/j.devcel.2019.10.027

Browning B. L., X. Tian, Y. Zhou, and S. R. Browning, 2021 Fast two-stage phasing of large-scale sequence data. Am. J. Hum. Genet. 108: 1880–1890. 10.1016/j.ajhg.2021.08.005

Cai X., C. Gao, H. Song, N. Yang, Q. Fu, et al., 2019 Characterization, expression profiling and functional characterization of cathepsin Z (CTSZ) in turbot (Scophthalmus maximus L.). Fish Shellfish Immunol. 84: 599–608. 10.1016/j.fsi.2018.10.046

Chang C. C., C. C. Chow, L. C. Tellier, S. Vattikuti, S. M. Purcell, et al., 2015 Second-generation PLINK: rising to the challenge of larger and richer datasets. Gigascience 4: s13742–015–0047–8. 10.1186/s13742-015-0047-8

Chen S., Y. Zhou, Y. Chen, and J. Gu, 2018 fastp: an ultra-fast all-in-one FASTQ preprocessor. Bioinformatics 34: i884–i890. 10.1093/bioinformatics/bty560

Delaneau O., H. Ongen, A. A. Brown, A. Fort, N. I. Panousis, et al., 2017 A complete tool set for molecular QTL discovery and analysis. Nat. Commun. 8: 15452. 10.1038/ncomms15452

Evans D. H., P. M. Piermarini, and K. P. Choe, 2005 The multifunctional fish gill: dominant site of gas exchange, osmoregulation, acid-base regulation, and excretion of nitrogenous waste. Physiol. Rev. 85: 97–177. 10.1152/physrev.00050.2003

Gjerde B. R., S. A. Boison, D. Hazlerigg, T. Ytrestøyl, T. Mørkøre, et al., 2025 Impact of three light smoltification regimes on performance and genetic parameters of traits in Atlantic salmon. Frontiers in Animal Science.

Grønvold L., M. J. van Dalum, A. Striberny, D. Manousi, T. Ytrestøyl, et al., 2024 Transcriptomic profiling of gill biopsies to define predictive markers for seawater survival in farmed Atlantic salmon. J. Fish Biol. 10.1111/jfb.16025

Iversen M., T. Mulugeta, B. Gellein Blikeng, A. C. West, E. H. Jørgensen, et al., 2020 RNA profiling identifies novel, photoperiod-history dependent markers associated with enhanced saltwater performance in juvenile Atlantic salmon. PLoS ONE 15: e0227496. 10.1371/journal.pone.0227496

Khaw H. L., B. Gjerde, S. A. Boison, E. Hjelle, and G. F. Difford, 2021 Quantitative genetics of smoltification status at the time of seawater transfer in atlantic salmon (salmo salar). Front. Genet. 12: 696893. 10.3389/fgene.2021.696893

Laneve P., F. Altieri, M. E. Fiori, A. Scaloni, I. Bozzoni, et al., 2003 Purification, cloning, and characterization of XendoU, a novel endoribonuclease involved in processing of intron-encoded small nucleolar RNAs in Xenopus laevis. J. Biol. Chem. 278: 13026–13032. 10.1074/jbc.M211937200

Li H., and R. Durbin, 2009 Fast and accurate short read alignment with Burrows-Wheeler transform. Bioinformatics 25: 1754–1760. 10.1093/bioinformatics/btp324

Maehr T., T. Wang, J. L. González Vecino, S. Wadsworth, and C. J. Secombes, 2012 Cloning and expression analysis of the transforming growth factor-beta receptors type 1 and 2 in the rainbow trout Oncorhynchus mykiss. Dev. Comp. Immunol. 37: 115–126. 10.1016/j.dci.2011.10.006

Malard F., K. Dias, M. Baudy, S. Thore, B. Vialet, et al., 2025 Molecular basis for the calcium-dependent activation of the ribonuclease EndoU. Nat. Commun. 16: 3110. 10.1038/s41467-025-58462-6

McCormick S. D., 2013 Smolt Physiology and Endochrinology, in Euryhaline Fishes; Fish Physiology,.

Morro B., P. Balseiro, A. Albalat, C. Pedrosa, S. Mackenzie, et al., 2019 Effects of different photoperiod regimes on the smoltification and seawater adaptation of seawater-farmed rainbow trout (Oncorhynchus mykiss): Insights from Na+, K+–ATPase activity and transcription of osmoregulation and growth regulation genes. Aquaculture 507: 282–292. 10.1016/j.aquaculture.2019.04.039

Panoutsopoulos A. A., A. H. De Crescenzo, A. Lee, A. M. Lu, A. P. Ross, et al., 2020 Pak1ip1 Loss-of-Function Leads to Cell Cycle Arrest, Loss of Neural Crest Cells, and Craniofacial Abnormalities. Front. Cell Dev. Biol. 8: 510063. 10.3389/fcell.2020.510063

Patro R., G. Duggal, M. I. Love, R. A. Irizarry, and C. Kingsford, 2017 Salmon provides fast and bias-aware quantification of transcript expression. Nat. Methods 14: 417–419. 10.1038/nmeth.4197

Renganaath K., and F. W. Albert, 2025 Trans-eQTL hotspots shape complex traits by modulating cellular states. Cell Genomics 5: 100873. 10.1016/j.xgen.2025.100873

Schwarz D. S., and M. D. Blower, 2014 The calcium-dependent ribonuclease XendoU promotes ER network formation through local RNA degradation. J. Cell Biol. 207: 41–57. 10.1083/jcb.201406037

Solbakken V. A., T. Hansen, and S. O. Stefansson, 1994 Effects of photoperiod and temperature on growth and parr-smolt transformation in Atlantic salmon (Salmo salar L.) and subsequent performance in seawater. Aquaculture 121: 13–27. 10.1016/0044-8486(94)90004-3

Soneson C., M. I. Love, and M. D. Robinson, 2015 Differential analyses for RNA-seq: transcript-level estimates improve gene-level inferences. [version 2; peer review: 2 approved]. F1000Res. 4: 1521. 10.12688/f1000research.7563.2

Srivastava A., L. Malik, H. Sarkar, M. Zakeri, F. Almodaresi, et al., 2020 Alignment and mapping methodology influence transcript abundance estimation. Genome Biol. 21: 239. 10.1186/s13059-020-02151-8

Strand J. E. T., D. Hazlerigg, and E. H. Jørgensen, 2018 Photoperiod revisited: is there a critical day length for triggering a complete parr-smolt transformation in Atlantic salmon Salmo salar? J. Fish Biol. 93: 440–448. 10.1111/jfb.13760

Takvam M., K. Sundell, H. Sundh, N. Gharbi, H. Kryvi, et al., 2024 New wine in old bottles: Modification of the Na^+^ /K^+^ -ATPase enzyme activity assay and its application in salmonid aquaculture. Rev. Aquacult. 16: 1087–1098. 10.1111/raq.12887

Tao J., Y. Chen, X. Bian, T. Cai, C. Song, et al., 2025 Prognostic and immunological implications of cathepsin Z overexpression in prostate cancer. Front. Immunol. 16: 1618487. 10.3389/fimmu.2025.1618487

Tsukazaki T., T. A. Chiang, A. F. Davison, L. Attisano, and J. L. Wrana, 1998 SARA, a FYVE domain protein that recruits Smad2 to the TGFbeta receptor. Cell 95: 779–791. 10.1016/s0092-8674(00)81701-8

Van der Auwera G. A., M. O. Carneiro, C. Hartl, R. Poplin, G. Del Angel, et al., 2013 From FastQ data to high confidence variant calls: the Genome Analysis Toolkit best practices pipeline. Curr. Protoc. Bioinformatics 11: 11.10.1-11.10.33. 10.1002/0471250953.bi1110s43

West A. C., Y. Mizoro, S. H. Wood, L. M. Ince, M. Iversen, et al., 2021 Immunologic profiling of the atlantic salmon gill by single nuclei transcriptomics. Front. Immunol. 12: 669889. 10.3389/fimmu.2021.669889

Ytrestøyl T., E. Hjelle, J. Kolarevic, H. Takle, A. Rebl, et al., 2023 Photoperiod in recirculation aquaculture systems and timing of seawater transfer affect seawater growth performance of Atlantic salmon (Salmo salar). J. World Aquaculture Soc. 54: 73–95. 10.1111/jwas.12880

Yu W., Z. Qiu, N. Gao, L. Wang, H. Cui, et al., 2011 PAK1IP1, a ribosomal stress-induced nucleolar protein, regulates cell proliferation via the p53-MDM2 loop. Nucleic Acids Res. 39: 2234–2248. 10.1093/nar/gkq1117

Zhang Q., M. Geng, K. Li, H. Gao, X. Jiao, et al., 2023 TGF-β1 suppresses the T-cell response in teleost fish by initiating Smad3- and Foxp3-mediated transcriptional networks. J. Biol. Chem. 299: 102843. 10.1016/j.jbc.2022.102843

Zhang J., and H. Zhao, 2023 eQTL studies: from bulk tissues to single cells. J. Genet. Genomics 50: 925–933. 10.1016/j.jgg.2023.05.003

